# Nesting strategy shapes territorial aggression but not testosterone: a comparative approach in female and male birds

**DOI:** 10.1101/2020.12.19.423623

**Authors:** SE Lipshutz, KA Rosvall

## Abstract

Our understanding of the proximate and ultimate mechanisms shaping competitive phenotypes primarily stems from research on male-male competition for mates, even though female-female competition is also widespread. Obligate secondary cavity-nesting has evolved repeatedly across avian lineages, providing a useful comparative context to explore how competition over limited nest cavities shapes aggression and its underlying mechanisms across species. Although evidence from one or another cavity-nesting species suggests that territorial aggression is adaptive in both females and males, this has not yet been tested in a comparative framework. We tested the hypothesis that cavity-nesting generates more robust territorial aggression, in comparison to close relatives with less restrictive nesting strategies. Our focal species were two obligate secondary cavity-nesting species and two related species with more flexible nesting strategies in the same avian family: tree swallow (*Tachycineta bicolor)* vs. barn swallow (*Hirundo rustica*); Eastern bluebird (*Sialia sialis*) vs. American robin (*Turdus migratorius*). We assayed conspecific territorial aggression, and found that cavity-nesting species physically attacked a simulated intruder more often than their close relatives. This pattern held for both females and males. Because territorial aggression is often associated with elevated testosterone, we also hypothesized that cavity-nesting species would exhibit higher testosterone levels in circulation. However, cavity-nesting species did not have higher testosterone in circulation for either sex, despite some correlative evidence that testosterone is associated with higher rates of physical attack in female tree swallows. Our focus on a competitive context that is relevant to both sexes – competition over essential breeding resources – provides a useful comparative framework for co-consideration of proximate and ultimate drivers of reproductive competition in females and males.

## Introduction

How do competitive traits evolve? In male animals, ornaments, armaments, and intense aggressive behavior are thought to be primarily driven by mating competition; indeed, variation in competitive traits in males largely maps on to interspecific variation in sexual selection and mating systems (Bro-Jørgensen, 2007; Cooney et al., 2019; Emlen and Oring, 1977; Göran, 1998; Miles et al., 2018). For females, early hypotheses considered female competitive traits as byproducts of correlated selection on male traits (Darwin, 1871; Lande, 1980). The alternative hypothesis, that female-female competition directly shapes the evolution of competitive traits in females, has since received abundant evidence (Clutton-Brock, 2009; Hare and Simmons, 2019). Across the tree of life, females engage in social competition and receive fitness benefits (Boersma et al., 2020; Bro-Jørgensen, 2002; Krieg and Getty, 2020; Rosvall, 2011, 2008; Sandell, 1998; Slagsvold and Lifjeld, 1994; Stockley and Bro-Jørgensen, 2011; Wu et al., 2018). It is no longer in question that intrasexual competition among females is adaptive. However, debate remains as to how exactly selection shapes the evolution of competitive traits in females, in part because researchers often focus on different modes of selection in the two sexes (i.e. sexual vs. social selection, Cain and Rosvall, 2014; Carranza, 2009; Clutton-Brock, 2009; Price, 2015; Riebel et al., 2019; Tobias et al., 2012; West-Eberhard, 1983). In essence, we need a unified framework on the evolution of competitive traits that applies to both sexes (sensu Emlen and Oring 1977).

This issue also comes into focus in evolutionary endocrinology, which has grappled with a mechanistic framework for the evolution of testosterone in females. Thirty years ago, the challenge hypothesis provided such a framework in males (Wingfield et al., 1990). One key component of this hypothesis is that interspecific variation in testosterone secretion is shaped by territorial aggression and trade-offs with parental care. Subsequent work has revealed higher levels of testosterone in species with enhanced territorial aggression or stronger degree of mating competition among males, at least in some vertebrate taxa (Garamszegi et al., 2005; Hirschenhauser et al., 2003; Mank, 2007; Marler and Trainor, 2020), but see (Goymann et al., 2019; Husak and Lovern, 2014; Moore et al., 2020). Research on the relationship between testosterone and female competition has accumulated more recently (Goymann and Wingfield, 2014; Ketterson et al., 2005; Rosvall et al., 2020). Despite the fact that testosterone is a natural and important part of female physiology (Drummond, 2006; Staub and De Beer, 1997), females tend to have lower levels of testosterone in circulation than males (Adkins-Regan, 2005), potentially due to sex-specific constraints on egg production and parental care, which could in turn influence male testosterone levels via intersexual coevolutionary processes (Ketterson et al., 2005). In females, macro-evolutionary patterns of testosterone are unrelated to mating system, degree of sexual dimorphism, and other metrics typically associated with male competition (Garamszegi, 2014; Goymann and Wingfield, 2014). Likewise, the correlative link between individual differences in testosterone and aggression is not well supported in females (reviewed in Rosvall et al., 2020), even in socially polyandrous species where females compete for mates (Lipshutz and Rosvall, 2020). However, exogenous testosterone may *experimentally* affect female aggression (Rosvall et al., 2020). In essence, there is some evidence linking testosterone to female aggression, but the application of hypotheses developed for males has not been fruitful in understanding how female aggression and its underlying mechanisms evolve (Rosvall et al., 2020, see also Duque-Wilckens and Trainor, 2017; Goymann and Wingfield, 2014). These observations highlight the need to test general principles on the evolution of competitive phenotypes using a framework that applies to both sexes.

One approach is to examine how behavior and physiology evolve when both sexes face strong selective pressures related to reproductive competition. We propose nesting strategy as a framework for the evolution of competitive phenotypes, because these resources are required for reproduction, and species vary in the flexibility vs. limited nature of this resource. A particularly limiting strategy is obligate secondary cavity-nesting, for which individuals must acquire natural or abandoned holes in order to nest and rear offspring (Newton, 1994; Nilsson, 1984; Zarnowitz and Manuwal, 1985). Cavity availability can be severely restricted, especially for obligate secondary cavity nesters who have no reproductive alternatives if they cannot maintain a pre-made cavity (Bunnell, 2013; Ibarra et al., 2017). Cavity-nesting has arisen independently at least 39 times in birds (Collias, 1997; Davidson et al., 2017), providing a useful comparative framework for testing behavioral and physiological hypotheses across multiple species. However, to our knowledge, there has not been a comparative evaluation of how cavity nesting shapes conspecific aggression, despite several lines of evidence that it should (elaborated below).

Here, we examine patterns of territorial aggression and testosterone secretion in 2 well studied obligate secondary cavity-nesting bird species in North America. We compare them each to a related species from the same avian family that has less restrictive nesting strategies, but otherwise similar life histories and ecologies. These non-cavity-nesting relatives utilize nest sites that are more broadly distributed in their environment, which presumably reduces selection to compete over nest sites. Focusing on the early spring period of territorial establishment, we hypothesize that competition for a limited cavity resource drives higher territorial aggression. This hypothesis stems from species-specific observations that aggressive competition over nesting cavities can escalate to injury or death (Duckworth, 2008; Leffelaar and Robertson, 1985), presumably because aggression has some adaptive value in obtaining or maintaining access to limited cavities (Albers et al., 2017; Duckworth and Badyaev, 2007; Krieg and Getty, 2018; Rosvall, 2008; Sandell and Smith, 1997; Szász et al., 2019). One prediction is that cavity nesters will have higher aggression and elevated testosterone secretion, regardless of sex. Alternatively, nesting strategy may better predict trait variation in one sex than the other. For instance, cavity-nesting may primarily differentiate female behavior, assuming that males (but not females) in non-cavity nesting species also experience strong competition for territories and/or mates. Or, testosterone may track nesting strategy, but only in males, if this hormone does not generate interspecific variation in female aggression. The specific constellation of outcomes will yield important insights into the evolution of competitive traits.

## Materials and Methods

### Study system

We focus on 4 extremely well-studied songbird species, with abundant information on their life history. In the Turdidae family, Eastern bluebirds (*Sialia sialis*) nest inside cavities (Gowaty and Plissner, 2020) and their relatives, American robins (*Turdus migratorius*), build their nests within a variety of trees and shrubs (Vanderhoff et al., 2020). In the Hirundinidae family, tree swallows (*Tachycineta bicolor*) nest inside cavities (Winkler et al., 2020), and their relatives, barn swallows (*Hirundo rustica*), build their nests inside barns (Brown and Brown, 2020). Except for nesting strategy, each within-family species pair is ecologically similar with regard to mating system, parental care, foraging strategy, migration, etc., and evolutionary distances are roughly similar within each pair ~18-20 million years (Kumar et al., 2017).

We conducted fieldwork around Bloomington, IN (39.142 N, 86.602 W), Lexington, KY (38.104 N, 84.489 W) and Monticello, IL (40.028 N, 88.573 W). In many songbirds, aggression and testosterone tends to peak during the pre-breeding period of territorial establishment and mate acquisition prior to egg laying (George and Rosvall, 2018; Ketterson et al., 2005; Slagsvold and Lifjeld, 1994). Therefore, we collected data during the territorial establishment phase of breeding but before egg-laying, which ranged mid-March to early May from 2018 – 2020, depending on species-specific phenology. We monitored eBird and checked nest boxes daily as males and females arrived from migration, formed territorial pairs, and defended territories by displacing intruders. We determined breeding status based on the stage of nest development. For American robins, we observed pairs for several hours to determine territorial boundaries. Barn swallows reuse clay nests across years, so we monitored which nests were currently claimed as individuals flew in and out of the barn between foraging bouts. Though some criteria used to determine breeding stage vary in a species-specific manner, our observations collectively suggest that all individuals were within the territorial establishment phase. Methods were approved by Indiana University IACUC #18-004 and all relevant federal and state permits.

### Simulated territorial intrusion

We simulated territorial intrusion in males and females using randomized combinations of 3-4 male or female taxidermy mounts and 3-5 conspecific vocalizations. In total, we assayed aggression in 116 individuals; see Table 1 for sample sizes specific to species and sexes. We began trials with a conspecific vocal lure to ensure the focal individuals noticed the mount and waited 30 seconds before beginning the trial. Over the span of 5 minutes, we measured a suite of aggressive behaviors involved in territorial defense, including physical contact with the mount and average distance from the mount. We ultimately focused on physical contact as our key aggressive behavior, because overt expressions of aggression like physical attacks represent the culmination of competitive interactions. We counted the number of 5-second intervals that contained physical contact with the mount, totaling to a maximum attack score of 60. We also analyzed distance from the mount to confirm that all individuals were present and engaged with the simulated territorial intrusion, whether or not they were aggressively attacking. We did not evaluate other aggressive behaviors or signals of aggressive intent, such as flyovers or songs, because these traits may not be expressed uniformly between the sexes or between the two avian families.

**Table 1:** Sample sizes for territorial intrusion (aggression) and plasma collection (testosterone).

|  | Eastern Bluebird |  | American Robin |  | Tree Swallow |  | Barn Swallow |  |
| --- | --- | --- | --- | --- | --- | --- | --- | --- |
|  | F | M | F | M | F | M | F | M |
| Aggression | 17 | 20 | 9 | 11 | 14 | 24 | 10 | 11 |
| Testosterone | 15 | 15 | 7 | 12 | 14 | 8 | 13 | 14 |

### Capture techniques and plasma collection

Our goal was to evaluate baseline levels of testosterone in circulation, rather than testosterone’s response to social stimulation, particularly in light of evidence that testosterone rarely elevates following a social challenge birds (Goymann et al., 2019). We achieved this goal in three ways: 1) by sampling individuals without a simulated territorial intrusion, i.e. passive collection (n = 68). 2) by sampling individuals immediately after a short simulated territorial intrusion (n = 16), and 3) by sampling individuals several days after a simulated territorial intrusion (n = 14). Working under the assumption that any social elevation of testosterone takes at least 15 minutes and peaks closer to 30 or 45 minutes after initial activation of the HPG axis (Jawor et al., 2006; Rosvall et al., 2016), we aimed to sample blood within 15 min from the beginning of the territorial intrusion. Capture latencies in the immediate group averaged 16 min 5 sec ± 1 min 14 sec, from intrusion start to blood sampling. Within the immediate collection group, there was also no relationship between testosterone and latency to sampling (Pearson’s correlation: r = 0.10, p = 0.73). There were also no differences in testosterone levels between individuals that were collected passively, immediately post-STI, or delayed post-STI for females (ANOVA: F_2,45_ = 0.44, p = 0.66), nor for males (ANOVA: F_2,46_ = 1.31, p = 0.28). As a consequence, we combined these groups for further analysis. In sum, we measured testosterone in 98 individuals – see Table 1 for sample sizes specific to each species and sex.

We captured individuals using a mist-net or a nestbox trap. Most animals were terminally collected for an ongoing study on comparative neurogenomics. We used an anaesthetic overdose of isoflurane, followed by decapitation and collection of trunk blood. We confirmed breeding status by examining whether females had small white follicles, and males had enlarged, white testes. Due to initially lower sample sizes, we obtained non-terminal collection from the brachial vein for some additional barn swallows, and we found no significant differences in testosterone across the two sampling types for this species (t = −0.93, df = 21.56, p = 0.36). We collected whole blood into heparinized BD Microtainers (product #365965) or heparinized microcapillary tubes and stored on an ice pack until we separated plasma by centrifuging for 10 minutes at 10,000 rpm. We stored plasma at −20°C for later testosterone assays.

### Testosterone enzyme immunoassay

We extracted steroids from plasma samples using diethyl ether (3x extractions) and reconstituted in 250μL assay buffer. We measured testosterone using a High Sensitivity Testosterone Enzyme Immuno- Assay kit (Enzo #ADI-900-176, Farmingdale, NY, USA) following methods described in George and Rosvall (2018). We used 50μL plasma from females, and 10μL from males. We calculated T concentration by comparing sample absorbance with the absorbance of the assay’s standard curve (Gen5 curve-fitting software, Biotek EPOCH plate reader, Winooski, VT, USA). Samples from 3 females and 2 males initially showed greater than 80% maximum binding, so we re-ran these samples at 20μL plasma to obtain values in the most sensitive part of the curve. Samples from 12 males initially showed less than 20% maximum binding, so we re-ran these samples with 10μL plasma reconstituted in 500μL assay buffer. In the case of plasma volume being insufficient (i.e. 18-19μL instead of 20 μL for two males, and 42μL instead of 50μL for one female), we added the remaining of water and calculated testosterone concentration accordingly; these samples still fell in the most sensitive part of the curve. We ran all samples in duplicate (duplicate CV = 4.0% ± 0.45). Each plate contained 3 duplicates of an extracted plasma pool, used to calculate the within and among plates variability. Intra-assay CV was 6.9% ± 0.36 and inter-plate CV was 13.3%. We suspect that this elevated inter-plate CV stems from the use of 3 different kit lots across the 3 years of this study, though we note that we balanced species and sexes within a plate each year.

### Statistical analyses

We conducted all statistics in R version 3.3.2 (R-Core-Team, 2016). We examined normality using a Shapiro-Wilk test, visualized distributions with histograms, and examined outliers using a Grubbs test in the R package ‘outliers’.

To evaluate whether nesting strategy predicted territorial aggression, our model included physical attacks as the response variable, with nest type, sex, and their interaction as fixed effects, and family as a random effect. Our dataset included many individuals that did not attack, resulting in excess zeros, so we ran a GLMM with zero-inflated Poisson distribution using the ‘mixed_model’ function in the GLMMadaptive package (Pinheiro and Bates, 1995). We used AICc in the MuMIn package (Barton, 2020) to compare this model with another using a Poisson distribution without zero-inflation, which did not perform as well (Δ AIC = 468.98). We used the package ‘performance’ to obtain R2 for variance from fixed effects (marginal R2) or fixed plus random effects (conditional R2) (Nakagawa et al., 2017).

To test whether species differed in their engagement with the simulated intrusion, regardless of the degree of their physical attack response, we compared average distance from the taxidermy mount. Our model included average distance from the mount as the response variable, with nest type, sex, and their interaction as fixed effects, and family as a random effect. Average distance was normally distributed, so we ran a LMM with the ‘lmer’ function in the lme4 package (Bates et al., 2015).

As a secondary goal, we evaluated sex differences in territorial aggression within each species. For each species, we compared males and females in their rate of physical attacks using Wilcoxon-tests. We also used Wilcoxon-tests to evaluate whether aggression was influenced by the sex of the taxidermy mount, i.e. whether females more aggressive towards female intruders and vice versa.

To compare baseline levels of testosterone in circulation across species, we ran a LMM with testosterone as the response variable, nest type, sex, and their interaction as fixed effects, and family as a random effect. We normalized testosterone level using a log scale transformation.

For a subset of tree swallows (n = 7 females, n = 5 males), we were able to assess variation in aggression and circulating testosterone for the same individuals. We used spearman correlations to evaluate the relationship between % of time attacking the mount and baseline testosterone in circulation.

## Results

### Cavity-nesting species had higher territorial aggression

Nesting strategy significantly predicted the number of physical attacks, such that cavity-nesting species attacked the taxidermy mount significantly more (β = 3.82, SE = 1.36, z = 2.82, p = 0.0049; overall model: conditional R^2^ = 0.97, marginal R^2^ = 0.97; Fig. 1). Sex was not a significant predictor of physical attack (β = 1.07, SE = 1.31, z = 0.81, p = 0.42), nor was the interaction between nesting strategy and sex (β = −1.21, SE = 1.31, z = −0.92, p = 0.36), indicating that higher aggression in cavity-nesting species was similar for both females and males.

**Figure 1:**
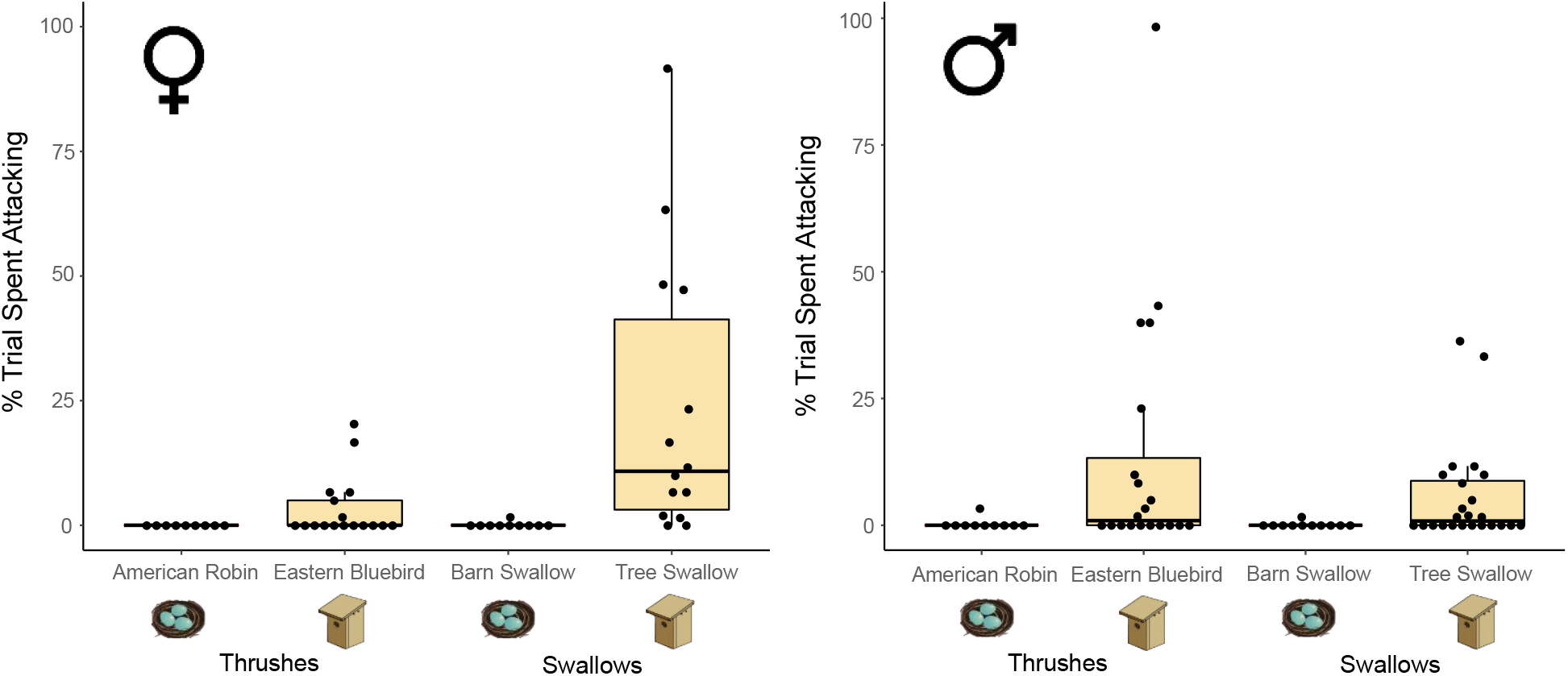
Boxplots show median and interquartile ranges for percent of time females (left panel) males (right panel) spent physically attacking taxidermy mount during simulated territorial intrusion in cavity-nesting species (light beige) and non-cavity-nesting species (dark seafoam).

We did not find a significant difference in average distance from the taxidermy mount, either based on nesting strategy, (β = −1.69, SE = 1.86, t = −0.91, p = 0.36; overall model: conditional R^2^ = 0.12, marginal R^2^ = 0.03; Fig. 2), sex (β = 0.98, SE = 2.02, t = 0.49, p = 0.63) nor the interaction between the two (β = 1.50, SE = 2.51, t = 0.60, p = 0.55). This suggests that mount placement was a salient stimulus, in that individuals from all species were responding to the simulated intruder in one way or another, regardless of whether the taxidermy mount was at a nest box.

**Figure 2:**
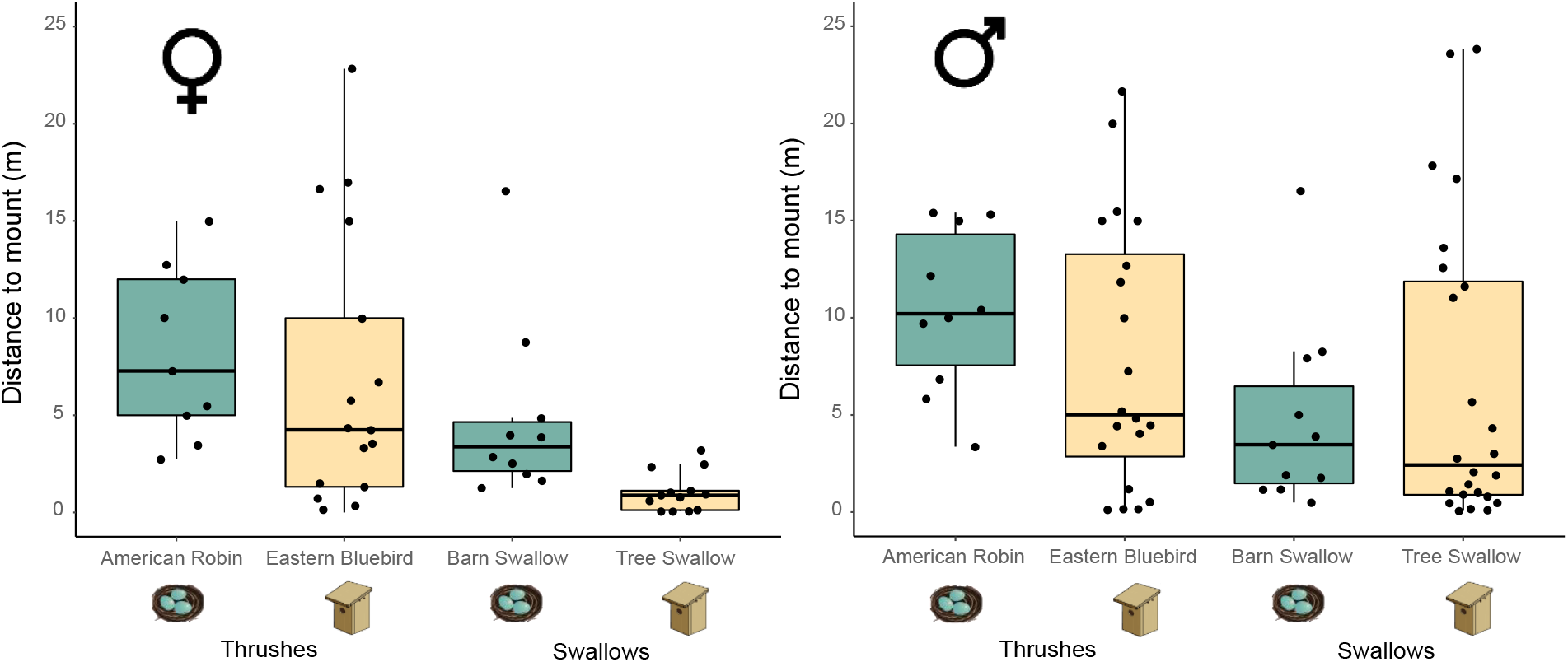
Boxplots show median and interquartile ranges for distance from taxidermy mount in females (left panel) and males (right panel) during simulated territorial intrusions in cavity-nesting species (light beige) and non-cavity-nesting species (dark seafoam).

### Sex differences in territorial aggression

For most species females and males attacked at similar rates, including for Eastern bluebirds (W = 133.5, p = 0.22), American robins (W = 40.5, p = 0.40), and barn swallows (W = 55.5, p = 1). However, in tree swallows, females attacked significantly more than males (W = 235.5, p = 0.0082). Territorial aggression was not related to the sex of the mount, except in Eastern bluebirds, males attacked male mounts significantly more than female mounts (W = 6.5, p = 0.008).

### Levels of testosterone did not relate to nesting strategy for either sex

Nesting strategy was not a significant predictor of testosterone in circulation (β = 0.019, SE = 0.11, t = 0.16, p = 0.87; conditional R^2^ = 0.66, marginal R^2^ = 0.65; Fig. 3). For all species, levels of testosterone in circulation were significantly higher in males than in females (β = 1.06, SE = 0.12, t = 9.13, p < 0.0001). There was no significant interaction between nesting strategy and sex (β = 0.026, SE = 0.16, t = 0.16, p = 0.87). Means and standard errors were: 0.19 ± 0.077 ng/mL and 2.99 ± 0.62 ng/mL for female and male robins, 0.22 ± 0.046 ng/mL and 2.83 ± 0.52 ng/mL for female and male bluebirds, 0.29 ± 0.054 ng/mL and 3.10 ± 0.80 ng/mL for female and male barn swallows, and 0.32 ± 0.099 ng/mL and 3.52 ± 1.03 ng/mL for female and male tree swallows.

**Figure 3:**
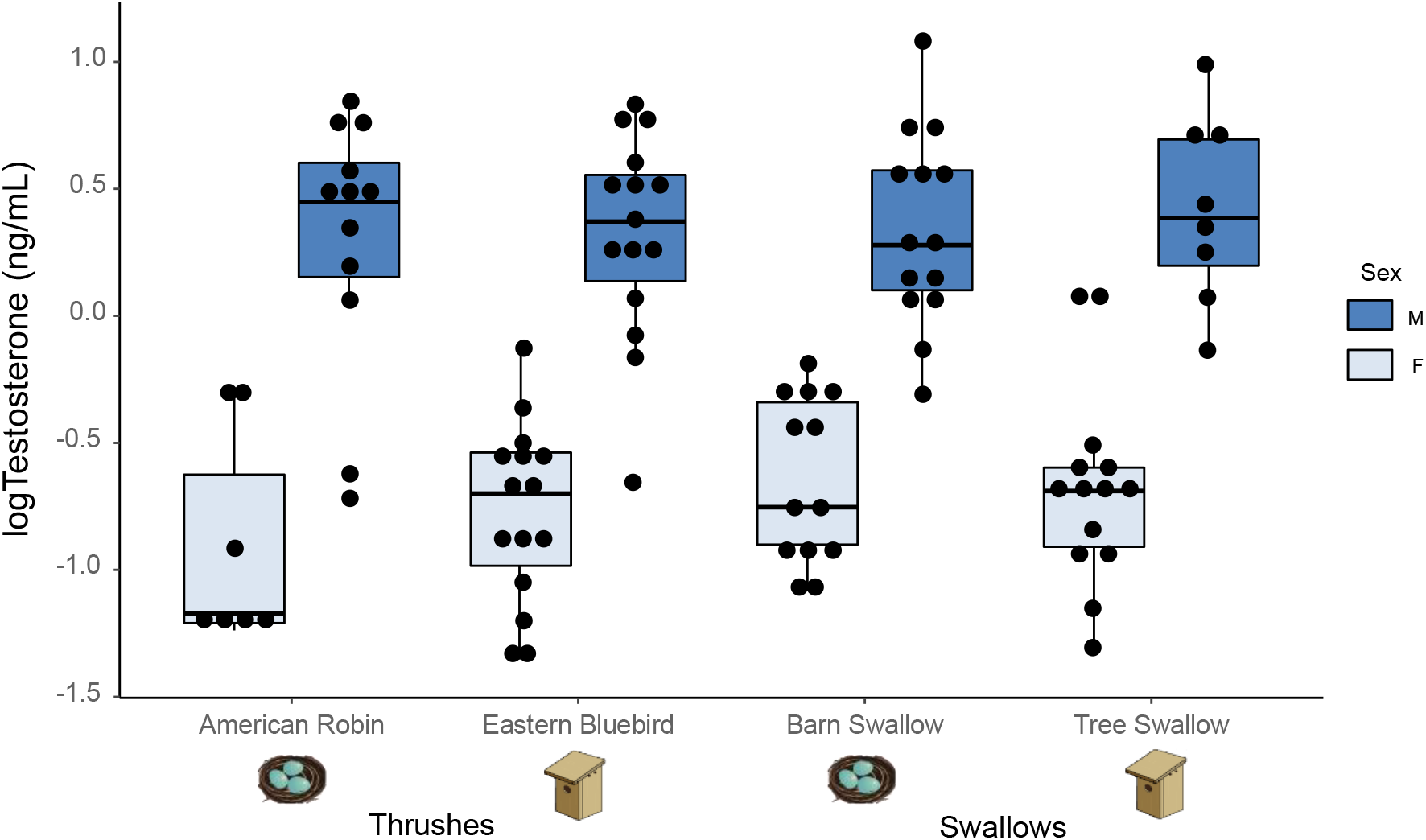
Levels of log testosterone (ng/mL) in circulation. Percent of time spent physically attacking taxidermy mount during simulated territorial intrusion. Females (light blue) males (dark blue). Cavity-nesters represented by nestbox, non-cavity-nesters represented by an open cup nest.

Tree swallows were the one species for which we had individually matched aggression and testosterone data for a subset of our study subjects. In females, the rate of physical attack was significantly, positively correlated with testosterone in circulation (rho = 0.96, p = 0.0028). There was no significant relationship between physical aggression and testosterone in male tree swallows (rho = −0.35, p = 0.56).

## Discussion

When it comes to the evolution of competitive traits, behavioral ecologists and endocrinologists have grappled with a functional and mechanistic framework that is broadly applicable to both females and males (Goymann and Wingfield, 2014; Ketterson et al., 2005; Lipshutz and Rosvall, 2020; Tobias et al., 2012), in part due to the separate consideration of selective pressures for each sex (Cain and Rosvall, 2014). In this comparative analysis, we propose and test a solution to this problem by evaluating the hypothesis that obligate secondary cavity nesting enhances territorial aggression and circulating testosterone levels in both sexes. This hypothesis was previously untested across species, though it lies at the core of species-specific studies on the adaptive value of competition for nesting cavities in birds (Duckworth, 2008; Gustafsson, 1988; Krieg and Getty, 2020; Leffelaar and Robertson, 1985; Rosvall, 2008). It also dovetails with evidence across other vertebrate taxa suggesting that females as well as males compete for limited breeding resources (Brandtmann et al., 1999; Hare and Simmons, 2019; Reedy et al., 2017; Stockley and Bro-Jørgensen, 2011; While et al., 2009; Wu et al., 2018), and that androgens may be involved (Cox et al., 2015; Davies et al., 2016; Desjardins et al., 2006; Woodley and Moore, 1999). We found that obligate secondary cavity-nesting is associated with higher conspecific aggression in two avian families, and this pattern applied to both females and males. Nesting strategy was not associated with higher levels of testosterone in circulation for either sex, despite some correlative evidence that female aggression is associated with higher testosterone in tree swallows. Our comparative approach focuses on reproductive competition for nesting sites, and in doing so, provides a useful framework for understanding proximate and ultimate drivers of competition in both sexes.

### Obligate cavity nesting and the evolution of aggression

Whereas most research on competitive traits focuses on male-male competition for mates or female-female competition for breeding resources, we demonstrate that interspecific variation in territorial aggression tracks nesting strategy in both sexes. In the Turdidae and Hirundinidae families, obligate secondary cavity-nesting species spent more of the simulated territorial intrusion physically attacking a conspecific mount compared to their relatives that have less restrictive nesting strategies. Past work on one or another obligate cavity nesting bird species found that aggression is beneficial for obtaining and maintaining a nesting territory (Duckworth and Badyaev, 2007; Krieg and Getty, 2020; Rosvall, 2008; Sandell and Smith, 1997; Szász et al., 2019). Our study finds that this adaptive behavioral trait is enhanced in species that compete for cavities, suggesting that nesting strategy is a potentially unifying driver of competitive phenotypes. This result adds to prior studies indicating that cavity-nesting may influence the evolution of a number of traits, via effects from predation, light regimes, heterospecific competition, and sexual selection (Davidson et al., 2017; Drury et al., 2020; Miles et al., 2018). Comparative studies like ours progress towards bridging within-species microevolutionary processes of behavioral adaptation with larger-scale macroevolutionary patterns, which are needed for a full understanding of behavioral evolution (Price et al., 2011).

Intriguingly, we did not find a sex-by-nesting type interaction on aggression, meaning that competition for nesting cavities is not necessarily a stronger selective force in one sex than the other. Sex-specific selection is a commonly hypothesized driver of sexually dimorphic phenotypes (Bell and Zamudio, 2012; Janicke et al., 2016; Rubenstein and Lovette, 2009; Shultz and Burns, 2017; but see Price, 2015). Likewise, when selection acts similarly in the two sexes, sexual dimorphism is reduced or absent (West-Eberhard, 1983). Accordingly, we see mutual ornaments and monomorphic traits such as brightly colored plumage and complex songs in female and male tropical birds that defend year-round territories (Dale et al., 2015; Tobias et al., 2011), as well as dewlaps in female and male in anoles (Harrison and Poe, 2012). Of course, males and females often differ in the direction or magnitude of some selective pressures. For instance, male territorial defense facilitates access to mates and mate guarding (Harts et al., 2016), and these factors should apply comparably to the 4 species in our study, all of which form socially monogamous pairs. In cavity-nesting females, aggression may also serve a number of functions, including securing exclusive access to mates (Sandell, 1998; Slagsvold et al., 1999, 1993), defense from conspecific brood parasitism (Gowaty and Wagner, 1988), ovicide avoidance (Krieg and Getty, 2020), and anti-predator defense (Winkler, 1992). Our comparative case study demonstrates that cavity-nesting rises above these potentially sex- and species-specific factors as an especially salient predictor of variation in aggression.

We found that in tree swallows, physical aggression was significantly higher in females than in males, regardless of the sex of the mount. This result is consistent with several lines of evidence that competition may be fierce for these females. In most tree swallow populations, nest sites are under threat of usurpation by an abundance of female floaters (Stutchbury and Robertson, 1987, 1985). Whereas male tree swallows may find breeding success through extra-pair copulations (Lifjeld et al., 1993), obtaining and maintaining cavity access is the only route to reproductive success in females because conspecific brood parasitism is rare (Barber and Robertson, 1999; Lombardo, 1988). In our examination of sex-specific aggression we also found that in Eastern bluebirds, males attacked male mounts significantly more than female mounts. A previous study found that bluebirds attacked in a sex-specific manner (Gowaty and Wagner, 1988), but this pattern was strongest during egg laying, when the risk of conspecific brood parasitism and extra-pair matings are highest. These two sex-specific results demonstrate that other factors may explain additional behavioral variance, beyond that which we linked to the shared drive to compete for cavities.

### Does testosterone facilitate parallel evolution of behavior?

If aggressive behavior has evolved in response to competition for cavities, a key next question is: how was this achieved at a mechanistic level? Evolutionary endocrinology often turns to testosterone to address divergence in traits related to reproductive or mating competition (Cox et al., 2009; Hau, 2007; Ketterson et al., 2009; Rosvall et al., 2016). Indeed, the challenge hypothesis proposes that species with more intense territorial aggression should have higher testosterone secretion (Wingfield et al., 1990). Our application of this hypothesis to competition for nesting cavities found parallel increases in aggression in cavity-nesting species, but this was not associated with concomitantly higher baseline levels of testosterone in circulation. This null relationship is not a female-specific pattern; testosterone did not track interspecific variation in male aggression either, even though males were clearly in breeding condition based on large testes and testosterone levels in the expected range (3.06 ng/ml average for males; 0.26 ng/mL for females). Although more sampling at additional breeding stages (e.g. during female fertility) is required to fully reject these links between testosterone and competition for nesting territories, our results mirror recent reports that the challenge hypothesis is context-dependent (Wingfield et al., 2020, 2019). Whereas testosterone levels have been related to alternative reproductive tactics (Küpper et al., 2015; Oliveira et al., 2005), male-female interactions (Goymann et al., 2019) and parental investment within species (Goymann and Flores Dávila, 2017; McGlothlin et al., 2007; Rosvall, 2013), as well as mating system across species (Garamszegi et al., 2005; Hirschenhauser and Oliveira, 2006), our study adds to a collection of many others that failed to find a relationship between testosterone and competitive traits across species (Goymann and Wingfield, 2014; Husak and Lovern, 2014). Uniquely, our study applies this null relationship in an appropriate context for both sexes: reproductive competition over breeding resources is essential for both female and male cavity-nesting species, and it is entirely analogous to territorial competition as laid out in the challenge hypothesis. This comparative framework better positions us to reject the hypothesis that that interspecific variation in testosterone tracks variation in aggression. This issue is especially critical in females, for which the association between testosterone and competition has rarely been assessed across species, let alone in a context related explicitly to female competition.

If interspecific variation in territorial aggression is not generated by differences in testosterone secretion at baseline levels, then is it time to leave testosterone behind? Mixed results at different levels of analysis suggest that this approach may be premature. For instance, we found a positive correlation between baseline testosterone and territorial aggression in female tree swallows, the species for which we had the most complete sampling of aggression and testosterone in the same individual. Correlational studies in other species have found mixed evidence for such a relationship (reviewed in Kempenaers et al., 2008; Rosvall et al., 2020; Williams, 2008) but there is good evidence that exogenous testosterone increases aggression in females (Rosvall et al., 2020), including tree swallows (Rosvall, 2013). There is still much to understand about why correlational links between testosterone and competitive traits *within* a population differ from patterns that emerge over larger evolutionary scales (Lipshutz et al., 2019). These mismatches may stem from context-dependent processes like local adaptation and phenotypic plasticity that generate or erode functional variation at one scale and not another (Agrawal, 2020; Hau and Goymann, 2015; Wingfield et al., 1997). Exploration of these processes in evolutionary endocrinology is promising (Cox, 2020; Lema, 2020), and more comparative approaches are needed. Levels of testosterone in circulation must also be considered within the entire sex steroid signaling system, which provides a diversity of routes to increase aggression without a change in testosterone (Ball and Balthazart, 2019; de Bournonville et al., 2020; Fuxjager and Schuppe, 2018; Schuppe and Fuxjager, 2019). Furthermore, aggression is regulated by many other mechanisms beyond testosterone, including arginine vasotosin, vasoactive intestinal peptide, serotonin, and progesterone (Goodson, 2005; Goodson et al., 2012; Goymann et al., 2008; Lischinsky and Lin, 2020; Nelson and Chiavegatto, 2001). As evolutionary behavioral endocrinology continues to embrace complexity, application of this comparative framework to more species will unveil the shared vs. diverse neuroendocrine mechanisms that facilitate behavioral evolution.

## Acknowledgments

This material is based upon work supported by the National Science Foundation Postdoctoral Research Fellowship in Biology under grant No. 1907134, NSF CAREER grant No. 1942192, Indiana Academy of Science, and Indiana University Department of Biology. Any opinion, findings, and conclusions or recommendations expressed in this material are those of the authors(s) and do not necessarily reflect the views of the National Science Foundation. We thank E. George, A. Bentz, S. Wolf, M. Woodruff, K. Content, L. Aguilar, S. Torneo, J. Kuske, T. Empson, M. Hauber, and M. Ward for assistance in the lab and field as well as feedback on the manuscript. We thank the IU Research and Teaching Preserve, Indiana Department of Natural Resources, D. Westneat, D. Dilcher, and Schaffer family for field site access. The authors declare no conflicts of interest.

## Literature Cited

Adkins-Regan, E., 2005. Hormones and Animal Social Behavior. Princeton University Press, Princeton, NJ.

Agrawal, A.A., 2020. A scale-dependent framework for trade-offs, syndromes, and specialization in organismal biology. Ecology 101, 1–24. https://doi.org/10.1002/ecy.2924

Albers, A.N., Jones, J.A., Siefferman, L., 2017. Behavioral differences among eastern bluebird populations could be a consequence of tree swallow presence: A pilot study. Front. Ecol. Evol. 5, 1–6. https://doi.org/10.3389/fevo.2017.00116

Ball, G.F., Balthazart, J., 2019. The neuroendocrine integration of environmental information, the regulation and action of testosterone and the challenge hypothesis. Horm. Behav. 104574. https://doi.org/10.1016/j.yhbeh.2019.104574

Barber, C.A., Robertson, R.J., 1999. Floater males engage in extrapair copulations with resident female tree swallows. Auk 116, 264–269. https://doi.org/10.2307/4089478

Barton, K., 2020. MuMIn: Multi‐model inference. R package version 1.43.17 75.

Bates, D., Maechler, M., Bolker, B.M., Walker, S., 2015. Fitting linear mixed-effects models using {lme4}. J. Stat. Softw. 67, 1–48. https://doi.org/10.18637/jss.v067.i01

Bell, R.C., Zamudio, K.R., 2012. Sexual dichromatism in frogs: Natural selection, sexual selection and unexpected diversity. Proc. R. Soc. B Biol. Sci. 279, 4687–4693. https://doi.org/10.1098/rspb.2012.1609

Boersma, J., Enbody, E.D., Jones, J.A., Nason, D., Lopez-Contreras, E., Karubian, J., Schwabl, H., 2020. Testosterone induces plumage ornamentation followed by enhanced territoriality in a female songbird. Behav. Ecol. 31, 1233–1241. https://doi.org/10.1093/beheco/araa077

Brandtmann, G., Scandura, M., Trillmich, F., 1999. Female-female conflict in the harem of a snail cichlid (Lamprologus ocellatus): Behavioural interactions and fitness consequences. Behaviour 136, 1123–1144. https://doi.org/10.1163/156853999501793

Bro-Jørgensen, J., 2007. The intensity of sexual selection predicts weapon size in male bovids. Evolution 61, 1316–1326. https://doi.org/10.1111/j.1558-5646.2007.00111.x

Bro-Jørgensen, J., 2002. Overt female mate competition and preference for central males in a lekking antelope. Proc. Natl. Acad. Sci. U. S. A. 99, 9290–9293. https://doi.org/10.1073/pnas.142125899

Brown, M.B., Brown, C.R., 2020. Barn Swallow (Hirundo rustica), in: Poole, A.F. (Ed.), The Birds of North America Online. Cornell Lab of Ornithology, Ithaca, NY. https://doi.org/10.2173/bna.452

Bunnell, F.L., 2013. Sustaining Cavity-Using Species: Patterns of Cavity Use and Implications to Forest Management. ISRN For. 457698.

Cain, K.E., Rosvall, K.A., 2014. Next steps for understanding the selective relevance of female-female competition. Front. Ecol. Evol. 2, 2012–2014. https://doi.org/10.3389/fevo.2014.00032

Carranza, J., 2009. Defining sexual selection as sex-dependent selection. Anim. Behav. 77, 749–751. https://doi.org/10.1016/j.anbehav.2008.11.001

Clutton-Brock, T., 2009. Sexual selection in females. Anim. Behav. 77, 3–11. https://doi.org/10.1016/j.anbehav.2008.08.026

Collias, N.E., 1997. On the origin and evolution of nest building by Passerine birds. Condor 99, 253–270.

Cooney, C.R., Varley, Z.K., Nouri, L.O., Thomas, G.H., Moody, C.J.A., Jardine, M.D., 2019. Sexual selection predicts the rate and direction of colour divergence in a large avian radiation. Nat. Commun. 10, 1773. https://doi.org/10.1038/s41467-019-09859-7

Cox, C.L., Hanninen, A.F., Reedy, A.M., Cox, R.M., 2015. Female anoles retain responsiveness to testosterone despite the evolution of androgen-mediated sexual dimorphism. Funct. Ecol. 29, 758–767. https://doi.org/10.1111/1365-2435.12383

Cox, R.M., 2020. Sex steroids as mediators of phenotypic integration, genetic correlations, and evolutionary transitions. Mol. Cell. Endocrinol. 502, 110668.https://doi.org/ https://doi.org/10.1016/j.mce.2019.110668

Cox, R.M., Stenquist, D.S., Calsbeek, R., 2009. Testosterone, growth and the evolution of sexual size dimorphism. J. Evol. Biol. 22, 1586–1598. https://doi.org/10.1111/j.1420-9101.2009.01772.x

Dale, J., Dey, C.J., Delhey, K., Kempenaers, B., Valcu, M., 2015. The effects of life history and sexual selection on male and female plumage colouration. Nature 527, 367–370. https://doi.org/10.1038/nature15509

Darwin, C., 1871. The descent of man and selection in relation to sex. Murray, London.

Davidson, G.L., Thornton, A., Clayton, N.S., Davidson, G.L., 2017. Evolution of iris colour in relation to cavity nesting and parental care in passerine birds. Biol. Lett. 13, 20160783.

Davies, C.S., Smyth, K.N., Greene, L.K., Walsh, D.A., Mitchell, J., Clutton-Brock, T., Drea, C.M., 2016. Exceptional endocrine profiles characterise the meerkat: Sex, status, and reproductive patterns. Sci. Rep. 6, 1–9. https://doi.org/10.1038/srep35492

de Bournonville, C., McGrath, A., Remage-Healey, L., 2020. Testosterone synthesis in the female songbird brain. Horm. Behav. 121, 104716. https://doi.org/10.1016/j.yhbeh.2020.104716

Desjardins, J.K., Hazelden, M.R., Van Der Kraak, G.J., Balshine, S., 2006. Male and female cooperatively breeding fish provide support for the “Challenge Hypothesis.” Behav. Ecol. 17, 149–154. https://doi.org/10.1093/beheco/arj018

Drummond, A.E., 2006. The role of steroids in follicular growth. Reprod. Biol. Endocrinol. 11, 1–11. https://doi.org/10.1186/1477-7827-4-16

Drury, J.P., Cowen, M.C., Grether, G.F., 2020. Competition and hybridization drive interspecific territoriality in birds. Proc. Natl. Acad. Sci. 117, 12923–12930. https://doi.org/10.1073/pnas.1921380117

Duckworth, R.A., 2008. Adaptive dispersal strategies and the dynamics of a range expansion. Am. Nat. 172, S4–S17. https://doi.org/10.1086/588289

Duckworth, R.A., Badyaev, A. V., 2007. Coupling of dispersal and aggression facilitates the rapid range expansion of a passerine bird. Proc. Natl. Acad. Sci. 104, 15017–15022. https://doi.org/10.1073/pnas.0706174104

Duque-Wilckens, N., Trainor, B.C., 2017. Behavioral neuroendocrinology of female aggression. https://doi.org/10.1093/acrefore/9780190264086.013.11

Emlen, S.T., Oring, L.W., 1977. Ecology, sexual selection, and evolution of mating systems. Science (80-.). 197, 215–223.

Fuxjager, M.J., Schuppe, E.R., 2018. Androgenic signaling systems and their role in behavioral evolution. J. Steroid Biochem. Mol. Biol. 184, 47–56. https://doi.org/10.1016/j.jsbmb.2018.06.004

Garamszegi, L.Z., 2014. Female peak testosterone levels in birds tell an evolutionary story: a comment on Goyman and Wingfield. Behav. Ecol. 25, 700–701. https://doi.org/10.1093/beheco/aru048

Garamszegi, L.Z., Eens, M., Hurtrez-Boussès, S., Møller, A.P., 2005. Testosterone, testes size, and mating success in birds: A comparative study. Horm. Behav. 47, 389–409. https://doi.org/10.1016/j.yhbeh.2004.11.008

George, E.M., Rosvall, K.A., 2018. Testosterone production and social environment vary with breeding stage in a competitive female songbird. Horm. Behav. 103, 28–35. https://doi.org/10.1016/j.yhbeh.2018.05.015

Goodson, J.L., 2005. The vertebrate social behavior network: Evolutionary themes and variations. Horm. Behav. 48, 11–22. https://doi.org/10.1016/j.yhbeh.2005.02.003

Goodson, J.L., Wilson, L.C., Schrock, S.E., 2012. To flock or fight: Neurochemical signatures of divergent life histories in sparrows. Proc. Natl. Acad. Sci. 109, 10685–10685.

Göran, A., 1998. Comparative evidence for the evolution of genitalia by sexual selection. Nature 393, 784–786.

Gowaty, P.A., Plissner, J.H., 2020. Eastern Bluebird (Sialia sialis), in: Poole, A.F. (Ed.), Birds of the World. Cornell Lab of Ornithology, Ithaca, NY. https://doi.org/10.2173/bow.easblu.01

Gowaty, P.A., Wagner, S.J., 1988. Breeding season aggression of female and male Eastern bluebirds (Sialia sialis) to models of potential conspecific and interspecific egg dumpers. Ethology 250, 238–250.

Goymann, W., Flores Dávila, P., 2017. Acute peaks of testosterone suppress paternal care: Evidence from individual hormonal reaction norms. Proc. R. Soc. B Biol. Sci. 284, 20170632. https://doi.org/10.1098/rspb.2017.0632

Goymann, W., Moore, I.T., Oliveira, R.F., 2019. Challenge hypothesis 2.0: A fresh look at an established idea. Bioscience 69, 432–442. https://doi.org/10.1093/biosci/biz041

Goymann, W., Wingfield, J.C., 2014. Male-to-female testosterone ratios, dimorphism, and life history − What does it really tell us? Behav. Ecol. 25, 685–699. https://doi.org/10.1093/beheco/aru019

Goymann, W., Wittenzellner, A., Schwabl, I., Makomba, M., 2008. Progesterone modulates aggression in sex-role reversed female African black coucals. Proc. R. Soc. B Biol. Sci. 275, 1053–1060. https://doi.org/10.1098/rspb.2007.1707

Gustafsson, L., 1988. Inter‐ and intraspecific competition for nest holes in a population of the Collared Flycatcher Ficedula albicollis. Ibis (Lond. 1859). 130, 11–16. https://doi.org/10.1111/j.1474-919X.1988.tb00951.x

Hare, R.M., Simmons, L.W., 2019. Sexual selection and its evolutionary consequences in female animals. Biol. Rev. 94, 929–956. https://doi.org/10.1111/brv.12484

Harrison, A., Poe, S., 2012. Evolution of an ornament, the dewlap, in females of the lizard genus Anolis. Biol. J. Linn. Soc. 106, 191–201. https://doi.org/10.1111/j.1095-8312.2012.01847.x

Harts, A.M.F., Booksmythe, I., Jennions, M.D., 2016. Mate guarding and frequent copulation in birds: A meta-analysis of their relationship to paternity and male phenotype. Evolution (N. Y). 70, 2789–2808. https://doi.org/10.1111/evo.13081

Hau, M., 2007. Regulation of male traits by testosterone: Implications for the evolution of vertebrate life histories. BioEssays 29, 133–144. https://doi.org/10.1002/bies.20524

Hau, M., Goymann, W., 2015. Endocrine mechanisms, behavioral phenotypes and plasticity: known relationships and open questions. Front. Zool. 12, 1–15.

Hirschenhauser, K., Oliveira, R.F., 2006. Social modulation of androgens in male vertebrates: Meta-analyses of the challenge hypothesis. Anim. Behav. 71, 265–277. https://doi.org/10.1016/j.anbehav.2005.04.014

Hirschenhauser, K., Winkler, H., Oliveira, R.F., 2003. Comparative analysis of male androgen responsiveness to social environment in birds: The effects of mating system and paternal incubation. Horm. Behav. 43, 508–519. https://doi.org/10.1016/S0018-506X(03)00027-8

Husak, J.F., Lovern, M.B., 2014. Variation in steroid hormone levels among Caribbean Anolis lizards: Endocrine system convergence? Horm. Behav. 65, 408–415. https://doi.org/10.1016/j.yhbeh.2014.03.006

Ibarra, J.T., Martin, M., Cockle, K.L., Martin, K., 2017. Maintaining ecosystem resilience: functional responses of tree cavity nesters to logging in temperate forests of the Americas. Sci. Rep. 7, 4467. https://doi.org/10.1038/s41598-017-04733-2

Janicke, T., Häderer, I.K., Lajeunesse, M.J., Anthes, N., 2016. Evolutionary Biology: Darwinian sex roles confirmed across the animal kingdom. Sci. Adv. 2, 1–10. https://doi.org/10.1126/sciadv.1500983

Jawor, J.M., Mcglothlin, J.W., Casto, J.M., Greives, T.J., Snajdr, E.A., Bentley, G.E., Ketterson, E.D., 2006. Seasonal and individual variation in response to GnRH challenge in male dark-eyed juncos (Junco hyemalis). Gen. c 149, 182–189. https://doi.org/10.1016/j.ygcen.2006.05.013

Kempenaers, B., Peters, A., Foerster, K., 2008. Sources of individual variation in plasma testosterone levels. Philos. Trans. R. Soc. B Biol. Sci. 363, 1711–1723. https://doi.org/10.1098/rstb.2007.0001

Ketterson, E.D., Atwell, J.W., McGlothlin, J.W., 2009. Phenotypic integration and independence: Hormones, performance, and response to environmental change. Integr. Comp. Biol. 49, 365–379. https://doi.org/10.1093/icb/icp057

Ketterson, E.D., Nolan, V., Sandell, M., 2005. Testosterone in females: mediator of adaptive traits, constraint on sexual dimorphism, or both? Am. Nat. 166, S85–S98. https://doi.org/10.1086/444602

Krieg, C.A., Getty, T., 2020. Fitness benefits to intrasexual aggression in female house wrens, Troglodytes aedon. Anim. Behav. 160, 79–90. https://doi.org/10.1016/j.anbehav.2019.12.001

Krieg, C.A., Getty, T., 2018. Female house wrens value the nest cavity more than exclusive access to males during conflicts with female intruders. Behaviour 155, 151–180. https://doi.org/https://doi.org/10.1163/1568539X-00003481

Kumar, S., Stecher, G., Suleski, M., Hedges, S.B., 2017. TimeTree: A Resource for Timelines, Timetrees, and Divergence Times. Mol. Biol. Evol. 34, 1812–1819. https://doi.org/10.1093/molbev/msx116

Küpper, C., Stocks, M., Risse, J.E., Dos Remedios, N., Farrell, L.L., McRae, S.B., Morgan, T.C., Karlionova, N., Pinchuk, P., Verkuil, Y.I., Kitaysky, A.S., Wingfield, J.C., Piersma, T., Zeng, K., Slate, J., Blaxter, M., Lank, D.B., Burke, T., 2015. A supergene determines highly divergent male reproductive morphs in the ruff. Nat. Genet. 48, 79–83. https://doi.org/10.1038/ng.3443

Lande, R., 1980. Sexual dimorphism, sexual selection, and adaptation in polygenic characters Author. Evolution (N. Y). 34, 292–305. https://doi.org/10.2307/2407393

Leffelaar, D., Robertson, R.J.., 1985. Nest usurpation and female competition for breeding opportunities by Tree Swallows. Wilson Bull. 97, 221–224.

Lema, S.C., 2020. The adaptive value of hormones: Endocrine systems as outcomes and initiators of evolution. Mol. Cell. Endocrinol. 517. https://doi.org/10.1016/j.mce.2020.110983

Lifjeld, J.T., Dunn, P.O., Robertson, R.J., Boag, P.T., 1993. Extra-pair paternity in monogamous tree swallows. Anim. Behav. https://doi.org/10.1006/anbe.1993.1028

Lipshutz, S.E., George, E.M., Bentz, A.B., Rosvall, K.A., 2019. Evaluating testosterone as a phenotypic integrator: From tissues to individuals to species. Mol. Cell. Endocrinol. 496, 110531. https://doi.org/10.1016/j.mce.2019.110531

Lipshutz, S.E., Rosvall, K.A., 2020. Neuroendocrinology of sex-role reversal. Integr. Comp. Biol. icaa046. https://doi.org/10.1093/icb/icaa046

Lischinsky, J.E., Lin, D., 2020. Neural mechanisms of aggression across species. Nat. Rev. Neurosci. 8, 536–546. https://doi.org/10.1038/nrn2174

Lombardo, M., 1988. Evidence of intraspecific brood parasitism in the Tree Swallow. Wilson Bull. 100, 126–128.

Mank, J.E., 2007. The evolution of sexually selected traits and antagonistic androgen expression in actinopterygiian fishes. Am. Nat. 169, 142–149. https://doi.org/10.1086/510103

Marler, C.A., Trainor, B.C., 2020. The challenge hypothesis revisited: Focus on reproductive experience and neural mechanisms. Horm. Behav. 123, 104645. https://doi.org/10.1016/j.yhbeh.2019.104645

McGlothlin, J.W., Jawor, J.M., Ketterson, E.D., 2007. Natural Variation in a Testosterone‐Mediated Trade‐Off between Mating Effort and Parental Effort. Am. Nat. 170, 864–875. https://doi.org/10.1086/522838

Miles, M.C., Schuppe, E.R., Ligon IV, R.M., Fuxjager, M.J., 2018. Macroevolutionary patterning of woodpecker drums reveals how sexual selection elaborates signals under constraint. Proc. R. Soc. B Biol. Sci. 285, 20172628.

Moore, I.T., Hernandez, J., Goymann, W., 2020. Who rises to the challenge? Testing the Challenge Hypothesis in fish, amphibians, reptiles, and mammals. Horm. Behav. 123, 104537. https://doi.org/10.1016/j.yhbeh.2019.06.001

Nakagawa, S., Johnson, P.C.D., Schielzeth, H., 2017. The coefficient of determination R^2^ and intra-class correlation coefficient from generalized linear mixed-effects models revisited and expanded. J. R. Soc. Interface 14. https://doi.org/10.1098/rsif.2017.0213

Nelson, R.J., Chiavegatto, S., 2001. Molecular basis of aggression. Trends Neurosci 24, 713–719. https://doi.org/10.1016/S0166-2236(00)01996-2

Newton, I., 1994. The role of nest sites in limiting the numbers of hole-nesting birds : A review. Biol. Conserv. 70, 265–276. https://doi.org/10.1016/0006-3207(94)90172-4

Nilsson, S.G., 1984. The evolution of nest-site selection among hole-nesting birds: the importance of nest predation and competition. Ornis Scand. 15, 167–175.

Oliveira, R.F., Ros, A.F.H., Gonçalves, D.M., 2005. Intra-sexual variation in male reproduction in teleost fish: A comparative approach. Horm. Behav. 48, 430–439. https://doi.org/10.1016/j.yhbeh.2005.06.002

Pinheiro, J.C., Bates, D.M., 1995. Approximations to the Log-Likelihood Function in the Nonlinear Mixed-Effects Model. J. Comput. Graph. Stat. 4, 12–35. https://doi.org/10.1080/10618600.1995.10474663

Price, J.J., 2015. Rethinking our assumptions about the evolution of bird song and other sexually dimorphic signals. Front. Ecol. Evol. 3, 1–6. https://doi.org/10.3389/fevo.2015.00040

Price, J.J., Clapp, M.K., Omland, K.E., 2011. Where have all the trees gone? The declining use of phylogenies in animal behaviour journals. Anim. Behav. 81, 667–670. https://doi.org/10.1016/j.anbehav.2010.12.004

R-Core-Team, 2019. R: A language and environment for statistical computing.

Reedy, A.M., Pope, B.D., Kiriazis, N.M., Giordano, C.L., Sams, C.L., Warner, D.A., Cox, R.M., 2017. Female anoles display less but attack more quickly than males in response to territorial intrusions. Behav. Ecol. 28, 1323–1328. https://doi.org/10.1093/beheco/arx095

Riebel, K., Odom, K.J., Langmore, N.E., Hall, M.L., 2019. New insights from female bird song: Towards an integrated approach to studying male and female communication roles. Biol. Lett. 15, 1–7. https://doi.org/10.1098/rsbl.2019.0059

Rosvall, K.A., 2013. Life history trade-offs and behavioral sensitivity to testosterone: An experimental test when female aggression and maternal care co-occur. PLoS One 8. https://doi.org/10.1371/journal.pone.0054120

Rosvall, K.A., 2011. Intrasexual competition in females: Evidence for sexual selection? Behav. Ecol. https://doi.org/10.1093/beheco/arr106

Rosvall, K.A., 2008. Sexual selection on aggressiveness in females: evidence from an experimental test with tree swallows. Anim. Behav. 75, 1603–1610. https://doi.org/10.1016/j.anbehav.2007.09.038

Rosvall, K.A., Bentz, A.B., George, E.M., 2020. How research on female vertebrates contributes to an expanded challenge hypothesis. Horm. Behav. 123, 104565. https://doi.org/10.1016/j.yhbeh.2019.104565

Rosvall, K.A., Bergeon Burns, C.M., Jayaratna, S.P., Ketterson, E.D., 2016. Divergence along the gonadal steroidogenic pathway: Implications for hormone-mediated phenotypic evolution. Horm. Behav. 84, 1–8. https://doi.org/10.1016/j.yhbeh.2016.05.015

Rubenstein, D.R., Lovette, I.J., 2009. Reproductive skew and selection on female ornamentation in social species. Nature 462, 786–789. https://doi.org/10.1038/nature08614

Sandell, M.I., 1998. Female aggression and the maintenance of monogamy: Female behaviour predicts male mating status in European starlings. Proc. R. Soc. B Biol. Sci. 265, 1307–1311. https://doi.org/10.1098/rspb.1998.0434

Sandell, M.I., Smith, H.G., 1997. Female aggression in the European starling during the breeding season. Anim. Behav. 53, 13–23. https://doi.org/10.1006/anbe.1996.0274

Schuppe, E.R., Fuxjager, M.J., 2019. Phenotypic variation reveals sites of evolutionary constraint in the androgenic signaling pathway. Horm. Behav. 115, 104538. https://doi.org/10.1016/j.yhbeh.2019.06.002

Schuppe, E.R., Fuxjager, M.J., 2018. High-speed displays encoding motor skill trigger elevated territorial aggression in downy woodpeckers. Funct. Ecol. 32, 450–460. https://doi.org/10.1111/1365-2435.13010

Shultz, A.J., Burns, K.J., 2017. The role of sexual and natural selection in shaping patterns of sexual dichromatism in the largest family of songbirds (Aves: Thraupidae). Evolution (N. Y). 71, 1061–1074. https://doi.org/10.1111/evo.13196

Slagsvold, T., Dale, S., Lampe, H.M., 1999. Does female aggression prevent polygyny? An experiment with pied flycatchers (Ficedula hypoleuca). Behav. Ecol. Sociobiol. 45, 403–410. https://doi.org/10.1007/s002650050577

Slagsvold, T., Lifjeld, J.T., 1994. Polygyny in birds: the role of competition between females for male parental care. Am. Nat. 143, 59–94.

Slagsvold, T., Ornis, S., Scandinavian, S., Apr, N., 1993. Female-Female Aggression and Monogamy in Great Tits Parus major. Ornis Scand. 24, 155–158.

Staub, N.L., De Beer, M., 1997. The role of androgens in female vertebrates. Gen. Comp. Endocrinol. 108, 1–24. https://doi.org/10.1006/gcen.1997.6962

Stockley, P., Bro-Jørgensen, J., 2011. Female competition and its evolutionary consequences in mammals. Biol. Rev. 86, 341–366.

Stutchbury, B.J., Robertson, R.J., 1987. Behavioral tactics of subadult female floaters in the tree swallow. Behav. Ecol. Sociobiol. 20, 413–419. https://doi.org/10.1007/BF00302984

Stutchbury, B.J., Robertson, R.J., 1985. Floating Populations of Female Tree Swallows. Auk 102, 651–654. https://doi.org/10.1093/auk/102.3.651

Szász, E., Jablonszky, M., Krenhardt, K., Markó, G., Hegyi, G., Herényi, M., Laczi, M., Nagy, G., Rosivall, B., Szöllősi, E., Török, J., Garamszegi, L.Z., 2019. Male territorial aggression and fitness in collared flycatchers: a long-term study. Sci. Nat. 106, 1–11. https://doi.org/10.1007/s00114-019-1606-0

Tobias, J.A., Gamarra-Toledo, V., García-Olaechea, D., Pulgarín, P.C., Seddon, N., 2011. Year-round resource defence and the evolution of male and female song in suboscine birds: Social armaments are mutual ornaments. J. Evol. Biol. 24, 2118–2138. https://doi.org/10.1111/j.1420-9101.2011.02345.x

Tobias, J.A., Montgomerie, R., Lyon, B.E., 2012. The evolution of female ornaments and weaponry: social selection, sexual selection and ecological competition. Philos. Trans. R. Soc. B Biol. Sci. 367, 2274–2293. https://doi.org/10.1098/rstb.2011.0280

Vanderhoff, N., Pyle, P., Patten, M., Sallabanks, R., James, F., 2020. American Robin (Turdus migratorius), in: Poole, A.F. (Ed.), Birds of the World. Cornell Lab of Ornithology, Ithaca, NY.

West-Eberhard, M.J., 1983. Sexual selection, social competition, and speciation. Q. Rev. Biol. 58, 155–183. https://doi.org/10.1111/j.0014-3820.2001.tb00769.x

While, G.M., Sinn, D.L., Wapstra, E., 2009. Female aggression predicts mode of paternity acquisition in a social lizard. Proc. R. Soc. B Biol. Sci. 276, 2021–9. https://doi.org/10.1098/rspb.2008.1926

Wiebe, K.L., 2016. Interspecific competition for nests: Prior ownership trumps resource holding potential for Mountain Bluebird competing with Tree Swallow. Auk 133, 512–519. https://doi.org/10.1642/AUK-16-25.1

Williams, T.D., 2008. Individual variation in endocrine systems: moving beyond the ‘tyranny of the Golden Mean.’ Philos. Trans. R. Soc. B Biol. Sci. 363, 1687–1698. https://doi.org/10.1098/rstb.2007.0003

Wingfield, J.C., Goymann, W., Jalabert, C., Soma, K.K., 2020. Reprint of “Concepts derived from the Challenge Hypothesis.” Horm. Behav. 123. https://doi.org/10.1016/j.yhbeh.2020.104802

Wingfield, J.C., Hegner, R.E., Dufty, A.M., Ball, G.F., 1990. The “Challenge Hypothesis”: Theoretical implications for patterns of testosterone secretion, mating systems, and breeding strategies. Am. Nat. 136, 829–846.

Wingfield, J.C., Jacobs, J., Hillgarth, N., 1997. Ecological constraints and the evolution of hormone-behavior interrelationships. Ann. N. Y. Acad. Sci. 807, 22–41. https://doi.org/10.1111/j.1749-6632.1997.tb51911.x

Wingfield, J.C., Ramenofsky, M., Hegner, R.E., Ball, G.F., 2019. Whither the challenge hypothesis? Horm. Behav. 123, 104588. https://doi.org/10.1016/j.yhbeh.2019.104588

Winkler, D.W., 1992. Causes and consequences of variation in parental defense behavior by tree swallows. Condor 94, 502–520.

Winkler, D.W., Hallinger, K.K., Ardia, D.R., Robertson, R.J., Stutchbury, B.J., Cohen, R.R., 2020. Tree Swallow (Tachycineta bicolor), in: Poole, A.F. (Ed.), Birds of the World. Cornell Lab of Ornithology, Ithaca, NY. https://doi.org/10.2173/bna.treswa.02

Woodley, S., Moore, M., 1999. Female territorial aggression and steroid hormones in mountain spiny lizards. Anim. Behav. 57, 1083–1089. https://doi.org/10.1006/anbe.1998.1080

Wu, Y., Ramos, J.A., Qiu, X., Peters, R.A., Qi, Y., 2018. Female–female aggression functions in mate defence in an Asian agamid lizard. Anim. Behav. 135, 215–222. https://doi.org/10.1016/j.anbehav.2017.11.023

Zarnowitz, E.J., Manuwal, D.A., 1985. The effects of forest management on cavity nesting birds in north-western Washington. J. Wildl. Manage. 49, 255–263.

